# Temporal scaling of human scalp-recorded potentials

**DOI:** 10.1101/2020.12.11.421180

**Authors:** Cameron D. Hassall, Jack Harley, Nils Kolling, Laurence T. Hunt

## Abstract

Much of human behaviour is governed by common processes that unfold over varying timescales. Standard event-related potential analysis assumes *fixed-duration* responses relative to experimental events. However, recent single unit recordings in animals have revealed neural activity *scales* to span different durations during behaviours demanding flexible timing. Here, we employed a general linear modelling approach using a novel combination of fixed-duration and variable-duration regressors to unmix fixed-time and scaled-time components in human magneto/electroencephalography (M/EEG) data. We use this to reveal consistent temporal scaling of human scalp-recorded potentials across four independent EEG datasets, including interval perception, production, prediction and value-based decision making. Between-trial variation in the temporally scaled response predicts between-trial variation in subject reaction times, demonstrating the relevance of this temporally scaled signal for temporal variation in behaviour. Our results provide a general approach for studying flexibly timed behaviour in the human brain.

**Significance Statement:** Neural activity is traditionally thought to occur over fixed time scales. However, recent animal work has suggested that some neural responses occur over varying timescales. We extended this animal result to humans by detecting temporally scaled signals non-invasively at the scalp in four different tasks. Our results suggest that temporal scaling is an important feature of cognitive processes known to unfold over varying timescales.

---

Action and perception in the real world require flexible timing. We can walk quickly or slowly, recognize the same piece of music played at different tempos, and form temporal expectations over long and short intervals. In many cognitive tasks, reaction time variability is modelled in terms of internal evidence accumulation [1], whereby the same dynamical process unfolds at different speeds on different trials.

Flexible timing is critical in our lives, yet despite several decades of research [2–5] its neural correlates remain subject to extensive debate. Due to their high temporal resolution, magnetoencephalography and electroencephalography (M/EEG) have played a particularly prominent role in understanding the neural basis of timing [5–13], and the method typically used to analyze such data has been the event-related potential (ERP), which averages event-locked responses across multiple repetitions. For example, this approach has been used to identify the presence of a slow negative-going signal during timed intervals. This signal, called the contingent negative variation (CNV) [14], is thought to be timing related because its slope depends inversely on the duration of the timed interval [7,8,12].

Crucially, the ERP analysis strategy implicitly assumes that neural activity occurs at *fixed-time* latencies with respect to experimental events. However, it has recently been shown that brain activity at the level of individual neurons can be best explained by a *temporal scaling* model [15,16], in which activity is explained by a single response that is stretched or compressed according to the length of the produced interval. When monkeys are cued to produce intervals of different lengths, the temporal scaling model explains the majority of variance in neural responses from medial frontal cortex (MFC) single units [15]. This suggests that one mechanism by which flexible motor timing can be achieved is by adjusting the speed of a common neural process, a perspective readily viewed through the lens of dynamical systems theory [16]. Consistent with the broad role played by dynamical systems in a range of neural computations [17,18], recent studies in neural populations have revealed time-warping as a common property across many different population recordings and behavioural tasks [19]. For example, temporal scaling is also implicit in the neural correlates of evidence integration during sensory and value-based decision making [20] (which itself has also been proposed as a mechanism for time estimation in previous work [21]).

Successfully characterising scaled-time components in humans could open the door to studying the role of temporal scaling in more complex, hierarchical tasks such as music production or language perception, as well as in patient populations in which timing is impaired [22]. Yet it is currently unclear how temporal scaling of neural responses may manifest at the scalp (if at all) using non-invasive recording in humans. This is because of the fixed-time nature of the ERP analysis strategy. Again, one component of the ERP called the CNV has been found to ramp at different speeds for different temporal intervals [7,8,12], suggestive of temporal scaling. Crucially though, any scaled activity would appear mixed at the scalp with fixed-time components due to the *superposition problem* [23].

We therefore developed an approach to *unmix* scaled-time and fixed-time components in the EEG (Fig 1a). Our proposed method builds on recently developed least square regression-based approaches [24–29] that have proven useful in unmixing fixed-time components that overlap with one another, such as stimulus-related activity and response-related activity. To overcome the superposition problem, these approaches use a convolutional general linear model (GLM) to deconvolve neural responses that are potentially overlapping. Following this work, we estimate the fixed-time ERPs using a GLM in which the design matrix is filled with time-lagged ‘stick functions’ (a regressor which is valued 1 around the timepoint of interest, and 0 otherwise). Importantly, the stick functions can overlap in time to capture overlap in the underlying neural responses (Fig 1b), and the degree of fit to neural data can be improved by adding a regularisation penalty to the model estimation [29]. In situations without any overlap, the GLM would exactly return the conventional ERP.

**Fig 1.**
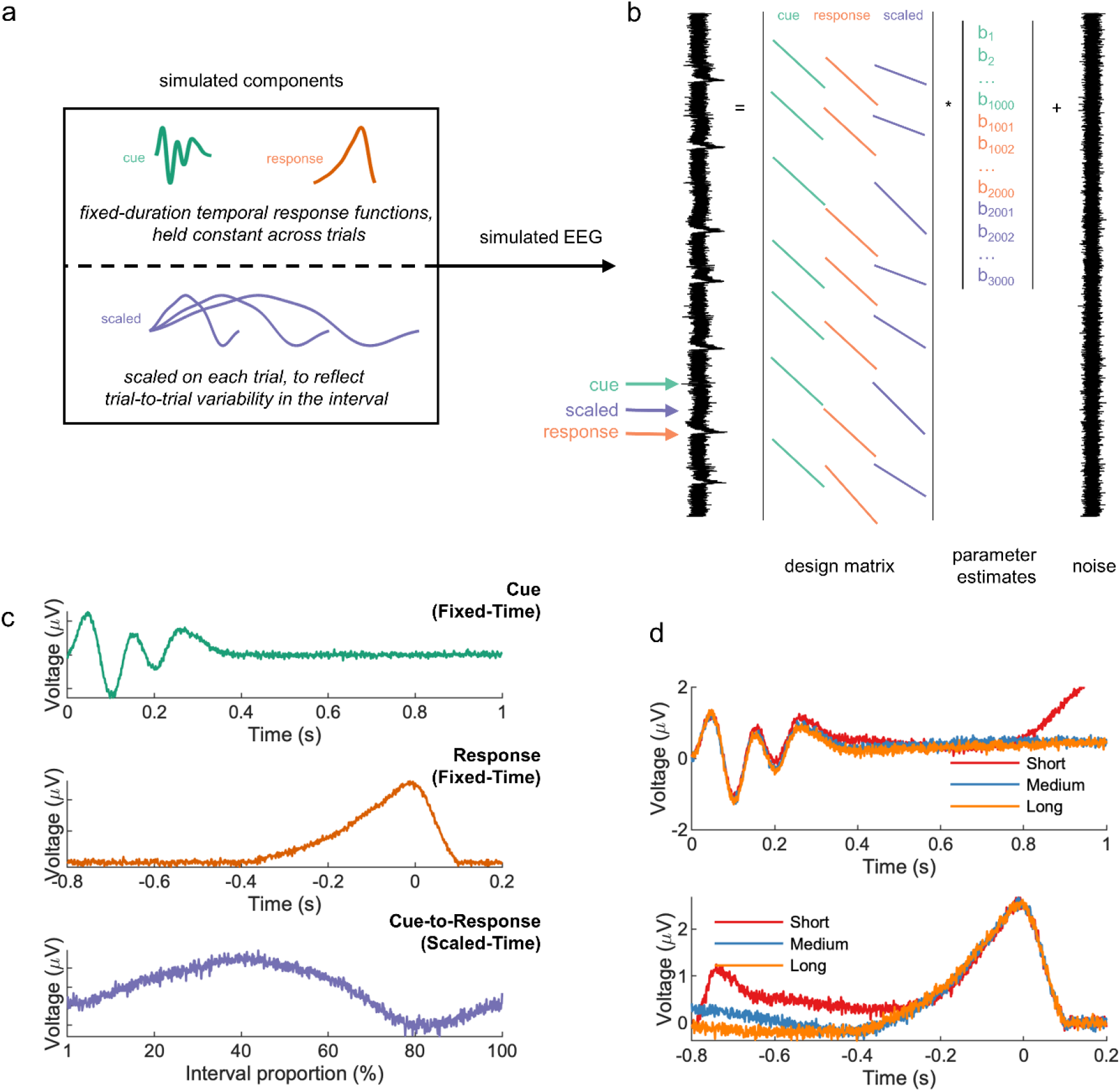
Regression based unmixing of simulated data successfully recovers scaled-time and fixed-time components. (a) EEG data were simulated by summing fixed-time components (cue and response), a scaled-time component with differing durations for different trials (short, medium, or long), and noise. (b) The simulated responses were unmixed via a GLM with stick basis functions: cue-locked, response-locked, and a single scaled-time basis spanning from cue to response (i.e., variable duration). (c) The GLM successfully recovered all three components, including the scaled-time component. (d) A conventional ERP analysis (cue-locked and response-locked averages) of the same data obscured the scaled-time component.

The key innovation that we introduce here is to allow for *variable-duration* regressors in such models, in addition to fixed-duration regressors, to test for the presence of scaled-time responses. In particular, we allow the duration of the stick function to vary depending upon the interval between stimulus and response, meaning that the same neural response can span different durations on different trials. Thus, rather than modelling the mean interval duration of each condition (e.g., via traditional ERPs), the proposed method captures trial-to-trial response variability. The returned scaled-time potential is no longer a function of real-world (‘wall clock’) time, but instead a function of the *percentage of time elapsed* between stimulus and response.

As a proof of concept, we simulated data at a single EEG sensor for an interval timing task, consisting of two *fixed-time components* (locked to cues and responses), and one *scaled-time component* spanning between cues and responses (Fig 1a). Our proposed method was successful in recovering all three components (Fig 1c), whereas a conventional ERP approach obscured the scaled-time component (Fig 1d). Crucially, in real EEG data we repeated this approach across all sensors, potentially revealing different scalp distributions (and hence different neural sources) for fixed-time versus scaled-time components.

By unmixing fixed and scaled components, our method goes beyond previous approaches for dealing with timing variability in EEG experiments. For example, the event-related timing of EEG trials can be aligned by translating either the entire waveform [30] or individual ERP components [31]. Such methods allow for component alignment, but do not involve any scaling. On the other hand, raw EEG can be scaled to align trials to a common time frame, e.g., through upsampling/downsampling [32] or dynamic time warping [33]. However, these methods are not designed to *unmix* fixed-time and scaled-time components. Finally, the effect of a continuous variable on EEG can be quantified using the method of temporal response functions (TRFs), another regression-based approach [27,34–36]. TRFs are particularly flexible in capturing different types of delay activity, e.g., during periods of growing expectancy [34] or while listening to fast/slow speech [35]. Our method is related to TRFs in that it involves the convolution of a to-be-estimated input signal with a continuous regressor. Unlike TRFs, however, the proposed method involves an additional scaling step in which the input signal is stretched or compressed (Fig 1, Supplementary Fig 1).

We also note that a time-frequency decomposition might also readily separate the responses at higher and lower-frequencies. Indeed, a wide range of neural oscillations have been implicated in time perception [37]. One might reasonably expect stretched/compressed signals to manifest differently in the time-frequency domain, e.g., as they correlate more strongly with different stretched/compressed versions of the same wavelet function. Unlike our proposed approach, however, a time-frequency decomposition is not readily designed to look for temporal scaling of the scaled-time response, namely the *same* neural response unfolding over *different* timescales on different trials. Nor will a time-frequency decomposition separate fixed-time responses from scaled-time responses if the signals occupy the same frequency band [38].

We used our approach to analyze EEG recorded across four independent datasets, comprising three interval timing tasks and one decision-making task. In the first task, participants produced a target interval (short, medium, or long) following a cue (Fig 2a). Feedback was provided, and participants were able to closely match the target intervals. In the second, participants evaluated a computer-produced interval (Fig 2b). The closer the produced interval was to the target interval, the more likely participants were to judge the response as ‘on time’. In the third (previously analyzed [39,40]) task, participants made temporal predictions about upcoming events based on rhythmic predictions (Fig 2c).

**Fig 2.**
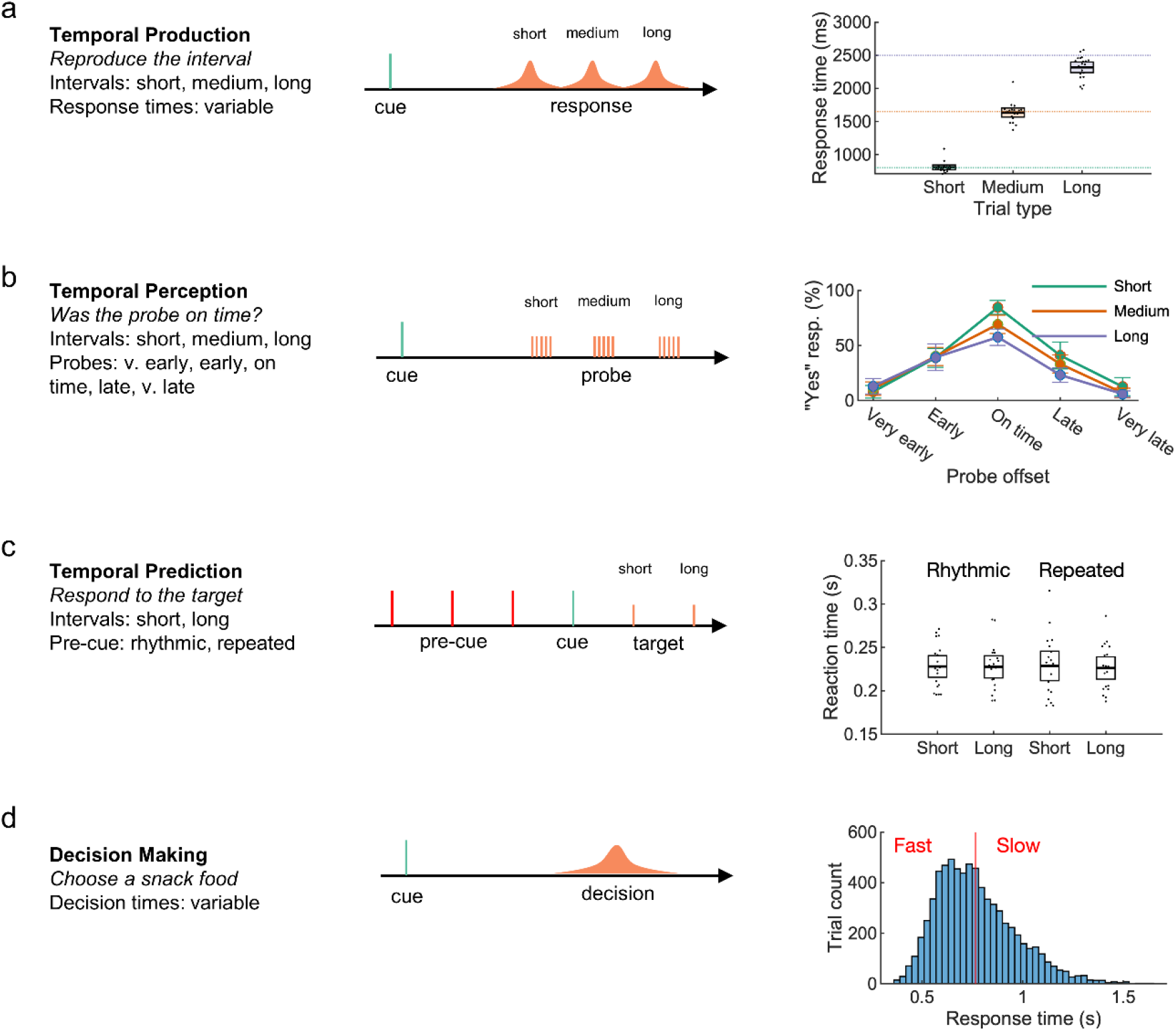
Datasets from three time-estimation and one decision-making paradigm were analyzed. In the temporal production task (a) participants successfully produced one of three cued intervals. In the temporal perception task (b) participants were able to properly judge a computer-produced interval. In a previously analyzed temporal prediction task[39,40], participants responded quickly to targets following either a rhythmic or repeated (non-rhythmic) cue. In a previously analyzed decision-making task[41,42] participants were cued to choose one of two snack food items, resulting in a range of response times (mean shown as red line). Error bars represent 95% confidence intervals.

In the fourth task (also previously analyzed [41,42]) participant chose between pairs of snack items (Fig 2d) – a process in which reaction time variability can be modelled as a process of internal evidence accumulation across time [43]. Neural activity related to evidence accumulation is measurable on the scalp as ramping activity that scales with decision difficulty. EEG for fast, easy trials increases at a faster rate compared to EEG for slow, difficult trials, indicating a higher rate of internal evidence accumulation [41]. Thus, we predicted that the EEG would contain an underlying scaled component associated with different rates of evidence accumulation.

In all four tasks, we observed a scaled-time component that was distinct from the preceding and following fixed-time components (Fig 3), which resembled conventional ERPs (Supplementary Fig 3). Typically, ERP components are defined by their polarity and scalp distribution [44]. The observed scaled-time components shared a common polarity (negative) and scalp distribution (central). In each task, cluster-based permutation testing revealed that the scaled-time component differed significantly from zero. The differences were driven by clusters spanning 36-87% in the production task (*p* < .001), 42-100% in the perception task (*p* < .001), 18-27% in the prediction task (*p* = .004), and 36-55% in the decision-making task (*p* < .001). For each task, including a scaled-time component improved model fit compared to a model with fixed-time components only: production: *t*(19) = −3.97, *p* < .001, Cohen’s *d* = − 0.89; perception: *t*(19) = −5.09, *p* < .001, Cohen’s *d* = −1.14; prediction: *t*(18) = −4.90, *p* < .001, Cohen’s *d* = −1.12; decision-making: *t*(17) = −7.77, *p* < .001, Cohen’s *d* = −1.83 (See Supplementary Table 8 for model errors). In many cases, scaled-time components were reliably observed at the single-subject level (Supplementary Fig 5-8).

**Fig 3.**
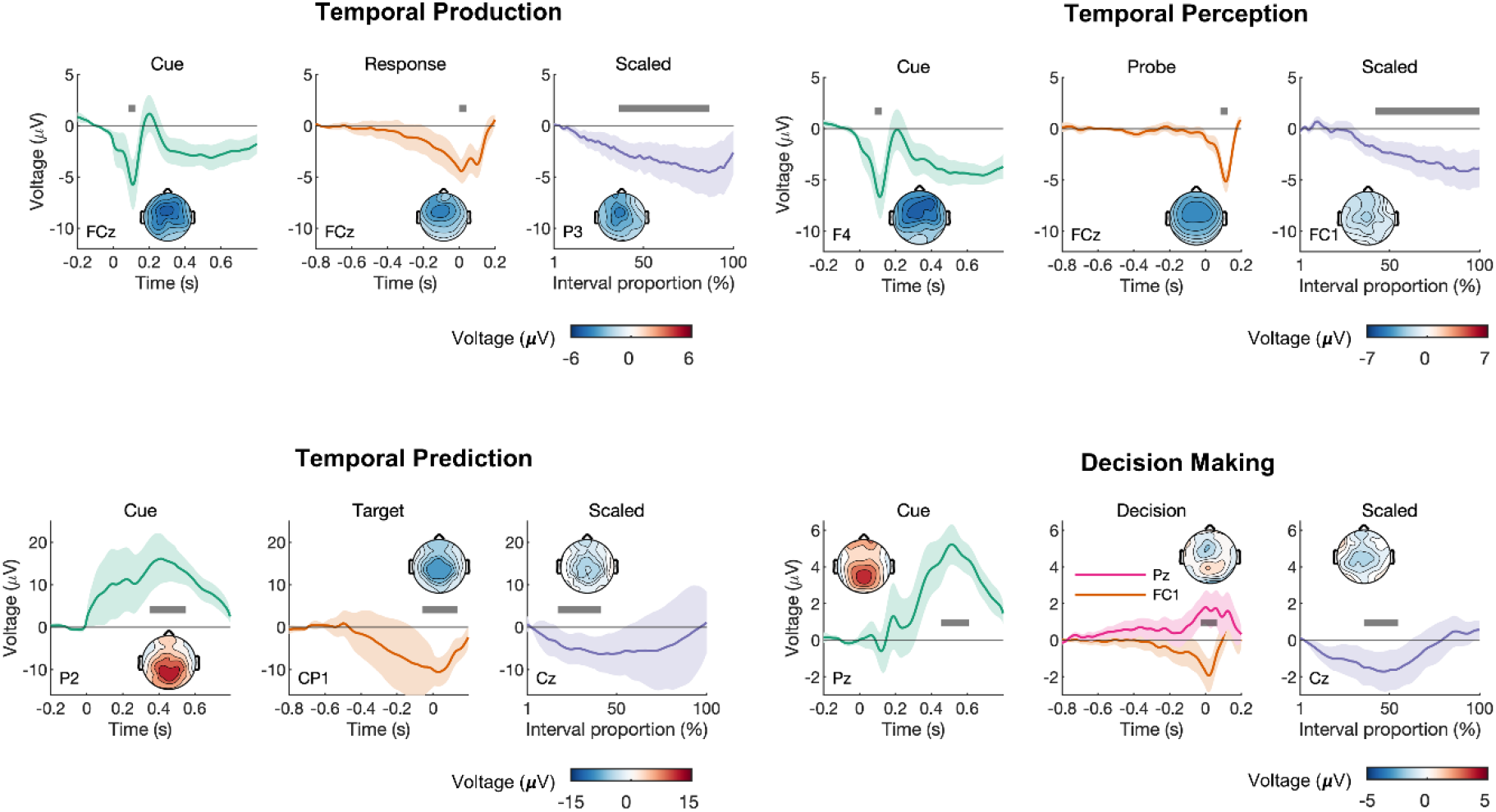
Scaled-time components were consistently observed across all four paradigms, with distinct scalp topographies from fixed-time components. Each had distinct fixed-time components relative to task-relevant events (left/middle columns), and a common negative scaled-time component over central electrodes, reflecting interval time (right column). The scalp topographies represent the mean voltage across the intervals indicated by the grey bars. For the fixed-time components, the intervals were chosen to visualize prominent deflections in the average waveform. For the scaled-time components, the intervals represent regions of significance as determined by cluster-based permutation tests. The error bars represent 95% confidence intervals.

To further validate our method, we quantified temporal scaling by computing a ‘scaling index’ [15] for each task and participant (Fig 4). To calculate this, we stretched/compressed each epoch to match the longest interval in each task, averaged by condition, then calculated the coefficient of determination for predicting the longer interval using stretched versions of the shorter intervals. We did this first on the raw data (‘Original’), then separately for the data containing only the fixed-time components (‘Fixed-only’, i.e., scaled-time components regressed out) and the scaled-time components (‘Scaled-only’, i.e., fixed-time components regressed out). In three out of four tasks, the scaling index for the scaled component exceeded the scaling index for the fixed component (production: *t*(19) = 2.95, *p* = .008, Cohen’s *d* = 0.66; perception: *t*(19) = 2.63, *p* = .017, Cohen’s *d* = 0.59; prediction: *t*(18) = 5.45, *p* < .001, Cohen’s *d* = 1.27; decision-making: *t*(17) = 5.45, *p* < .001, Cohen’s *d* = 1.29).

**Fig 4.**
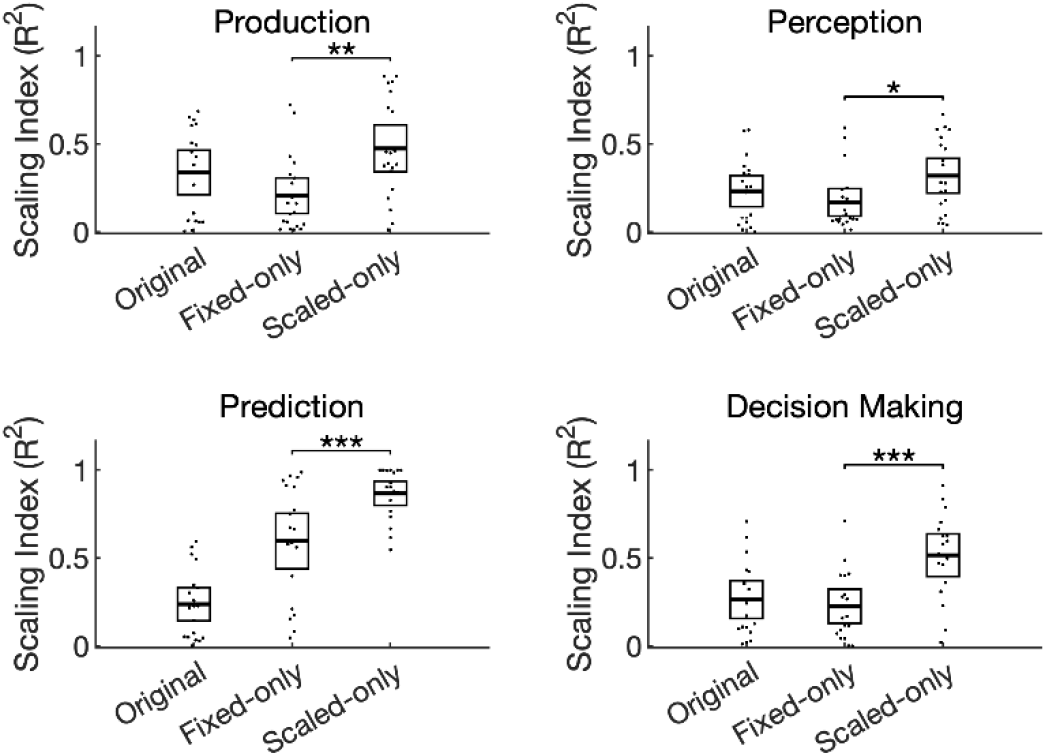
The unmixed signals differed quantitatively in their degree of scaling. The scaling index, defined as the coefficient of determination between epochs after stretching, was first computed for the raw data (‘Original’) and after isolating either the fixed (‘Fixed-only’) or scaled (‘Scaled-only’) components. In all four tasks the scaled-time components had a greater scaling index compared to the fixed time components. Dots represent individual participants and error bars represent 95% confidence intervals.

We then examined how the scaled-time component relates to behavioural variability: does the latency of the scaled-time component predict participants’ response time? We focussed on the temporal production and decision-making tasks, in which the interval duration was equal to the response time. As response time varied from trial to trial, so did the modelled scaled component. To measure component latency, we applied an approach developed in [45,46], using principal component analysis (PCA) to model delay activity over central electrodes in the temporal production task. The approach works by detecting latency shifts in a common underlying component [45,46]. Unlike simple peak detection, PCA can account for a range of waveform dynamics (e.g., multiple peaks). We first regressed out the fixed-time component as identified by the GLM, resulting in a dataset that consisted only of the residual scaled-time activity. We then computed the average scaled-time activity for each of the three interval conditions (Fig 5a-c). PCA was applied separately to each interval. This consistently revealed a first principal component that matched the shape of the scaled-time component and a second principal component that matched its *temporal derivative*. This analysis confirms the presence of the scaled-time component in our data, as it is the first principal component of the residuals after removing fixed-time components. Crucially, adding or subtracting the second principal component captures variation in the *latency* of the scaled-time component (Supplementary Fig 4). Across response time quantiles, we found that PC2 scores (Supplementary Table 6) were significantly related to response times (Fig 5d), *F*(2,38) = 6.18, *p* = .005, η*_p_*^2^ = 0.25, η*_g_*^2^ = 0.19). This implies that the earlier in time that the scaled-time component peaked, the faster the subject would respond on that trial. This result was replicated in the decision-making task, *F*(2,34) = 4.18, *p* = .02, η_*p*_^2^ = 0.20, η_*g*_^2^ = 0.18 (Fig 5e).

**Fig 5.**
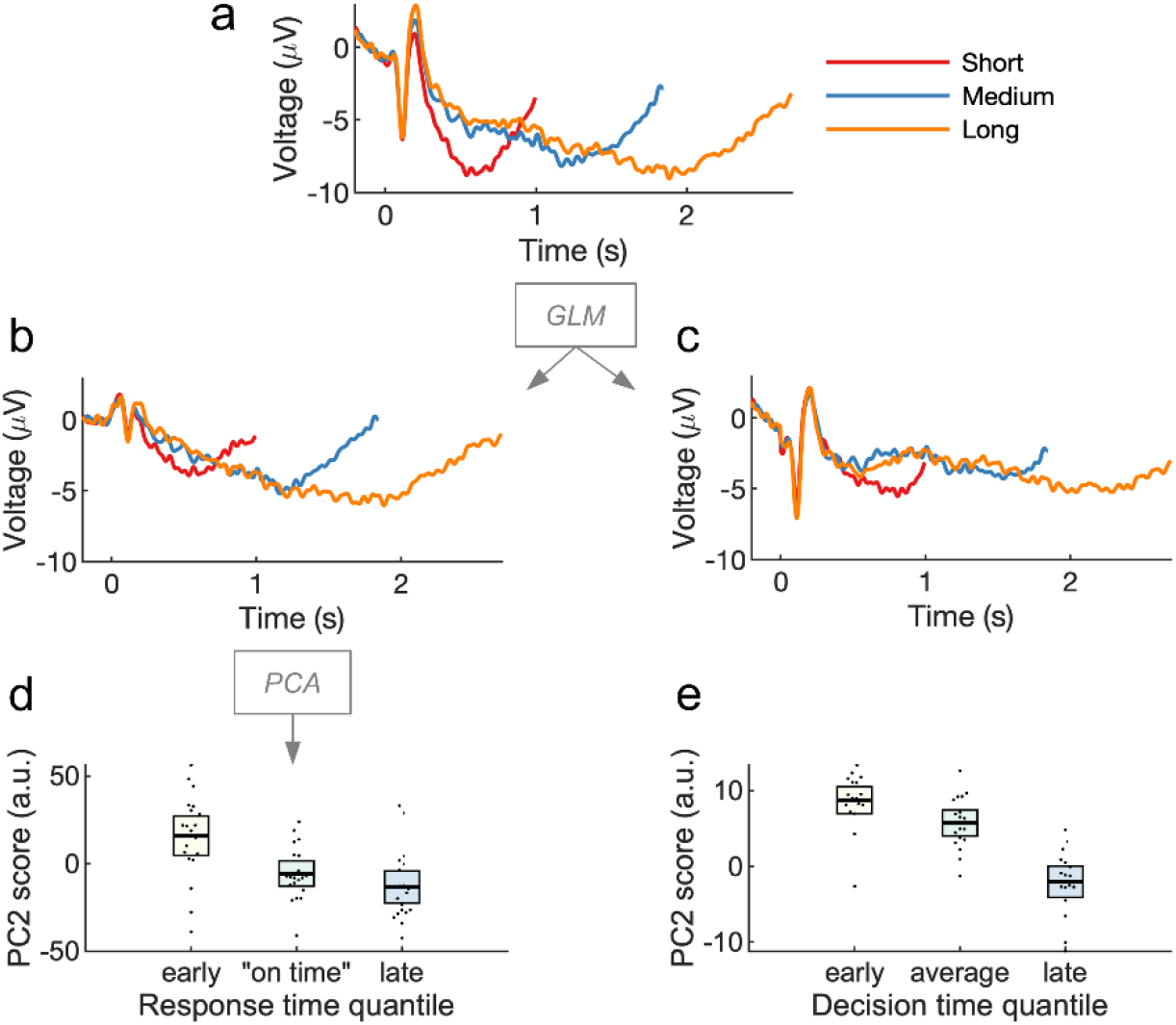
Variation in scaled-time components predicts behavioural variation in time estimation. Cue-locked EEG, shown as ERPs in (a), was analyzed via GLM. To visualize the unmixing of scaled-time and fixed-time components, the residual (noise) was recombined with either the scaled-time component (b) or the fixed-time components (c). PCA was run on the ‘scaled-time plus residual’ EEG. The second principal component resembled the temporal derivative or ‘rate’ of the scaled component (Supplementary Fig 5). (d) PC2 scores depended on response time, implying the scaled-time component peaked earlier for fast responses and later for slow responses. (e) The effect replicated in a decision-making task. Error bars represent 95% confidence intervals.

## Discussion

Our results provide a general method for recovering temporally scaled signals in human M/EEG, where scaled-time components are mixed at the scalp with conventional fixed-time ERPs. We focused here on tasks that have been widely used in the timing literature, namely interval production, perception, and prediction, as well as an example of a cognitive task that exhibits variable reaction times across trials (value-based decision making). Distinct scaled-time components and scalp topographies were revealed in all four tasks. These results suggest that flexible cognition relies on temporally scaled neural activity, as seen in recent animal work [15,16].

The existence of temporally scaled signals at the scalp may not be surprising to those familiar with the study of time perception. Because of its excellent temporal resolution, EEG has long been used to study delay activity in interval timing tasks. As discussed, one signal of interest has been a ramping frontal-central signal called the CNV, which we observed in our conventional ERP analysis (Supplementary Fig 3). Notably, CNV slope has been interpreted as an accumulation signal in pacemaker-accumulation models of timing [7,8,11,13]. Our work differs from these previous studies in one important respect. In a conventional ERP analysis, delay activity is assumed to occur over fixed latencies. The CNV is thus computed by averaging over many cue-locked EEG epochs of the same duration. In contrast, we have considered the possibility that scalped-recorded potentials reflect a mixture of both fixed-time and scaled-time components. By modelling fixed-time and scaled-time components separately, we revealed scaled activity that was common across all timed intervals. This, in turn, is consistent with a recent class of models of timing that propose time estimation reflects the variable speed over which an underlying dynamical system unfolds [16–18].

We also observed temporally scaled activity in a decision-making task, a somewhat surprising result given that the task did not have an explicit timing component (participants made simple binary decisions [41]). Nevertheless, time is the medium within which decisions are made [47]. Computationally, the timing of binary decisions can be captured in a drift diffusion model as the accumulation of evidence in favour of each alternative [1]. This accumulation is thought to be indexed by an ERP component called the central parietal positivity (CPP) [48]. There is evidence that the slope of the CPP – which can be either stimulus-locked or response-locked – captures the rate of evidence accumulation [49]. For faster/easier decisions the CPP climbs more rapidly compared to slower/harder decisions [48,49]. Perhaps these effects can also be explained by stretching/compressing a common scaled-time component while holding stimulus- and response-related activity constant. Furthermore, variation in the scaled-time component is relevant to decision making according to our results: it predicts *when* a decision will be made. However, we also note that the topography observed in our scaled-time component was a negative-going potential rather than positive (Fig 3d). This can potentially be explained by the standard CPP-like ERP [41] being a mixture of our observed negative scaled-time topography with the positive fixed-time topographies.

Although our approach makes no assumptions about the overall shape of the scaled-time component, it does assume a consistent, linear scaling across intervals. This is an assumption that could be relaxed in a more complex model, e.g., using spline regression [25]. We also note that although we made no a priori predictions about waveform shape, some between-task similarities and differences were noted in the resulting scaled-time components. For example, similar responses were seen in the tasks for which the interval of interest ended with a motor response (temporal production and decision making – see Fig 3a,d). In both cases, activity immediately preceding the response depended on a ramping, fixed-time, motor-related component, with little contribution from a scaled component. A similar observation was made in the temporal prediction task – activity just before the appearance of the target depended on anticipatory fixed-time activity, not scaled activity (Fig 3c). In contrast, pre-probe activity in the temporal perception task showed almost no fixed-time activity, but a robust scaled-time component (Fig 3b). The reason for this difference cannot be identified by the current experiments, however. First, the perception and prediction tasks involved different tasks instructions (‘listen for the probe’ versus ‘respond to the target’). Second, the probe/target distributions differed in the two tasks; the mean duration was 75% likely in the prediction task, but only 20% likely in the perception task. We therefore speculate that scaled activity may be somewhat task dependent.

Our approach is not only conceptually different from previous work that models variability in timing using a regression framework [27,34–36], it is also a mechanistically important finding. It indicates the brain may support flexible timing by adapting the duration of an otherwise consistent neural response. This can be understood as varying the rate of a dynamical system [17,18] during interval estimation. Although there is evidence for such temporally scaled responses in the monkey neurophysiology literature (e.g., [15,16], which inspired the current study), we are not aware of any direct evidence in support of this idea in humans. Indeed, it goes against the dominant framework of fixed-duration responses that has thus far dominated M/EEG analysis.

Although we have focused here on interval timing and decision-making tasks, we anticipate other temporally-scaled EEG and MEG signals will be discovered for cognitive processes known to unfold over varying timescales. For example, the neural basis of flexible (fast/slow) speech production and perception is an active area of research [50–52], and may involve a form of temporal scaling [32]. Flexible timing is also important across a vast array of decision-making tasks, where evidence accumulation can proceed quickly or slowly depending on the strength of the evidence [20]. Flexible timing helps facilitate a range of adaptive behaviours via temporal attention [4], while disordered timing characterizes several clinical disorders [53], underscoring the importance of characterising temporal scaling of neural responses in human participants.

## Acknowledgements

This research was funded by a Natural Sciences and Engineering Research Council of Canada (NSERC: https://www.nserc-crsng.gc.ca/) Postdoctoral Fellowship to C.D.H. (PDF 546078 – 2020), a BBSRC (https://bbsrc.ukri.org/) AFL Fellowship (BB/R010803/1) to N.K., a Sir Henry Dale Fellowship from the Royal Society (https://royalsociety.org) and Wellcome (https://wellcome.org/) to L.T.H. (208789/Z/17/Z), a NARSAD Young Investigator Award from the Brain and Behavior Research Foundation (https://www.bbrfoundation.org/) to L.T.H., and supported by the NIHR Oxford Health Biomedical Research Centre (https://oxfordhealthbrc.nihr.ac.uk/). The Wellcome Centre for Integrative Neuroimaging was supported by core funding from Wellcome Trust (203139/Z/16/Z). For the purpose of Open Access, the author has applied a CC BY public copyright licence to any Author Accepted Manuscript version arising from this submission.

## Author Contributions

N.K. and L.T.H. conceived the experiments and methodology. C.D.H. and L.T.H. designed the experiments and developed the methodology. C.D.H. and J.H. performed the experiments (except for the prediction task, where data was downloaded from a previous publication [39,40]). C.D.H. and L.T.H. analyzed the data. C.D.H. and L.T.H. wrote the manuscript with input from the other authors.

## Additional information

**Supplementary information** is available for this paper.

**Correspondence and requests for materials** should be addressed to C.D.H. or L.T.H.

## Methods

### Simulations

We simulated cue-related and response-related EEG in a temporal production task using MATLAB 2020a (Mathworks, Natick, USA). Cue and response were separated by either a short, medium, or long interval. During the delay period, we simulated a scaled response that stretched or compressed to fill the interval. All three responses (cue, response, scaled) were summed together at appropriate lags (short, medium, or long), with noise – see Fig 1a. In total, we simulated 50 trials of each condition (short, medium, long).

To unmix fixed-time and scaled-time components, we used a regression-based approach [24,25,54] in which the continuous EEG at one sensor *Y* is modelled as a linear combination of the underlying event-related responses β, which are unknown initially. The model can be written in equation form as:

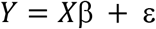

where *X* is the design matrix and ε is the residual EEG not accounted for by the model. *X* contains as many rows as EEG data points, and as many columns as predictors (that is, the number of points in the estimated event-related responses). In our case, *X* was populated by ‘stick functions’ – non-zero values around the time of the modelled events, and zeros otherwise. We included in *X* two fixed-time components, the cue and the response, as stick functions of fixed EEG duration (with variables set to 1). In other words, the height of the fixed-time stick function was constant across events of the same type and equal to its width. To model a temporally-scaled response, we used the MATLAB *imresize* function (Image Processing Toolbox, R2020b) with ‘box’ interpolation to stretch/compress a stick function so that it spanned the duration between cue and response (other interpolation methods were tried – see Supplementary Fig 1 – but this choice had little effect on the results). Thus, the duration of the scaled stick function varied from trial to trial (Fig 1b). The goal here was to estimate a single scaled-time response to account for EEG activity across multiple varying delay periods. For the fixed-time responses, each column of X represents a latency in ms before/after an experimental event; by contrast, for the scaled-time responses, each column of X represents the *percentage* of time that has elapsed between *two* events (stimulus and response). Simulation code is available at https://github.com/chassall/temporalscaling.

### Production and Perception Tasks

#### Participants

Participants completed both the production and perception tasks within the same recording session. We tested ten university-aged participants, 5 male, 2 left-handed, *M_age_* = 23.40, 95% CI [21.29, 25.51]. Participants had normal or corrected-to-normal vision and no known neurological impairments. This study was approved by the Medical Sciences Interdivisional Research Ethics Committee at the University of Oxford and participants provided informed consent. Following the experiment, participants were compensated £20 (£10 per hour of participation) plus a mean performance bonus of £3.23, 95% CI [2.92, 3.55].

#### Apparatus and Procedure

Participants were seated approximately 64 cm from a 27-inch LCD display (144 Hz, 1 ms response rate, 1920 by 1080 pixels, Acer XB270H, New Taipei City, Taiwan). Visual stimuli were presented using the Psychophysics Toolbox Extension [55,56] for MATLAB 2014b (Mathworks, Natick, USA). Participants were given written and verbal instructions to minimize head and eye movements. The goal of the production task was to produce a target interval and the goal of the perception task was to judge whether or not a computer-produced interval was correct.

The experiment was blocked with ten trials per block. There were 18 production blocks and 18 perception blocks, completed in random order. Prior to each block, participants listened to five isochronic tones indicating the target interval. Beeps were 400 Hz sine waves of duration 50 ms and an onset/offset ramping to a point 1/8 of the length of the wave (to avoid abrupt transitions). The target interval was either short (0.8 s), medium (1.65 s), or long (2.5 s).

In production trials, participants listened to a beep then waited the target time before responding. Feedback appeared after a 400-600 ms delay (uniform distribution) and remained on the display for 1000 ms. Feedback was a ‘quarter-to’ clockface to indicate ‘too early’, a ‘quarter-after’ clockface to indicate ‘too late’, or a checkmark to indicate an on-time response. Feedback itself was determined by where the participant’s response fell relative to a window around the target duration. The response window was initialized to +/− 100 ms around each target, then changed following each feedback via a staircase procedure: increased on each side by 10 ms following a correct response and decreased by 10 ms following an incorrect response (either too early or too late).

In perception trials, participants heard two beeps, then were asked to judge the correctness of the interval, that is, whether or not the test interval matched the target interval. Test intervals (very early, early, on time, late, very late) were set such that each subsequent interval was 25% longer than the previous (see Supplementary Table 1). Participants were then given feedback on their judgement – a checkmark for a correct judgement, or an ‘x’ for an incorrect judgement.

For each task, participants gained 2 points for each correct response and lost 1 point for each incorrect response. At the end of the experiment points were converted to a monetary bonus at a rate of £0.01 per point.

#### Data Collection

In the perception task we recorded participant response time from cue, trial outcome (early, late, on time), and staircase-response window. In the production task, we recorded trial ‘ on time’ judgements (yes/no), and trial outcome (correct/incorrect).

We recorded 36 channels of EEG, referenced to AFz. Data were recorded at 1000 Hz using a Synamps amplifier and CURRY 8 software (Compumetrics Neuroscan, Charlotte, USA). The electrodes were sintered Ag/AgCl (EasyCap, Herrsching, Germany). 31 of the electrodes were laid out according to the 10-20 system. Additional electrodes were placed on the left and right mastoids, on the outer canthi of the left and right eyes, and below the right eye. The reference electrode was placed at location AFz, and the ground electrode at Fpz.

### Prediction Task

In this previously published [39,40] experiment, 19 participants responded to the onset of a visual target following a visual warning cue. The delay between cue and target was either short (700 ms) or long (1300 ms) and, in some conditions, congruent with a preceding stimulus stream. Only congruent trials were included in the current analysis (i.e., the ‘valid’ trials in the ‘rhythmic’ and ‘repeated’ conditions). Each trial was preceded by a 500 ms fixation cross subtending 0.6°of visual angle. During the pre-cue period, participants were shown a flashing stimulus for 4-6 repetitions to indicate the target interval. The stimulus was a centrally presented black disc (1.2°) that appeared on the display for 100 ms. In the rhythmic condition the black disc appeared every 700 ms or 1300 ms (‘short’ or ‘long’). In the repeated condition, a red disc appeared either 700 ms or 1300 ms after the appearance of the black disc, followed by a variable delay period of either 1500-1900 ms (short) or 1900-2700 ms (long). Following the pre-cue period participants were then shown the warning cue, a white disc (1.2°) that appeared for 100 ms. After either a short or long delay (700 ms or 1300 ms) the target appeared – a green 1.2°disc – for 100 ms, followed by the participant’s response. The experimental program recorded the response time (time since the onset of the target). See Supplementary Fig 2c and [39,40] for more detail.

### Decision-Making Task

In this experiment, also previously published [41,42], 18 participants were presented with two snack foods and asked to pick one. This was not an interval timing task and on average participants took 763 ms, 95% CI [713, 813], to respond. Trials began with the appearance of a centrally presented fixation cross (0.6°) for 2-4 s followed by the presentation of the snack items (3°across, in total). Participants were asked to indicate their preference by making a left or right button press within a 1.25 s window. The experimental program recorded the response time (time since the onset of the snack items). See Supplementary Fig 2d and [41] for more detail.

### Data Analysis

#### Behavioural data

For the production task, we computed the mean produced interval for each participant. For the perception task, we computed mean likelihood of responding yes to the ‘on time’ prompt, for each condition (short, medium, long) and interval (very early, early, on time, late, very late). For the prediction task, we computed the mean reaction time for each analyzed condition (rhythmic, repeated) and interval (short, long). For the decision-making task, we computed the mean response (decision) time. See Fig 2 and Supplementary Tables 2-4 for behavioural results.

#### EEG Preprocessing

For all three timing tasks, EEG was preprocessed in MATLAB 2020b (Mathworks, Natick, USA) using EEGLAB [57]. We first down-sampled the EEG to 200 Hz, then applied a 0.1-20 Hz bandpass filter and 50 Hz notch filter. The EEG was then re-referenced to the average of the left and right mastoids (and AFz recovered in the production/perception tasks).

Ocular artifacts were removed using independent component analysis (ICA). The ICA was trained on 3-second epochs of data following the appearance of the fixation cross at the beginning of each trial. Ocular components were identified using the *iclabel* function and then removed from the continuous data.

EEG for the decision-making task was already preprocessed prior to our analysis. This was a simultaneous EEG-fMRI recording, and preprocessing included the removal of MR-related artifacts via filtering and principal component analysis, as well as a 0.5-40 Hz bandpass filter. In line with our other analyses, we re-referenced the EEG to the average of TP7 and TP8 (located close to the mastoids) and applied an additional 20 Hz low-pass filter.

#### ERPs

To construct conventional event-related potentials (ERPs), we first created epochs of EEG around cues (all tasks), responses (perception task), probes (production task), targets (prediction task), and decisions (decision-making task). Cue-locked ERPs extended from 200 ms pre-cue to either 800, 1650, or 2500 ms post-cue (the short, medium, and long targets) in the perception/production tasks, 700 or 1300 ms in the prediction task (the short and long targets), and 600 ms in the decision-making task. Epochs were baseline-corrected using a 200 ms pre-cue window. We also constructed epochs from 800, 1650, or 2500 ms prior to the response/probe in the production/perception tasks, 700 or 1300 ms prior to the target in the prediction task, and 600 prior to the decision in the decision-making task to 200 ms after the response/probe/target/decision. A baseline was defined around the event of interest (mean EEG from −20 to 20 ms) and removed in all cases except for the decision-making task, in line with the original analysis [41]. We then removed any trials in which the sample-to-sample voltage differed by more than 50 μV or the voltage change across the entire epoch exceeded 150 μV. We then created conditional cue and response/probe/target/decision averages for each participant and task: production/perception (short, medium, and long), prediction (short and long), and decision-making (early and late, via a median split [41]). Finally, participant averages in the timing tasks were combined into grand-average waveforms at electrode FCz, a location where timing-related activity has been previously observed [5] and Pz in the decision-making task, in line with the previously published analysis [41] (Supplementary Fig 3).

#### rERPs

To unmix fixed-time and scaled-time components in our EEG data, we estimated regression-ERPs (rERPs) following the same GLM procedure we used with our simulated data, but now applied to each sensor. We used a design matrix consisting of a regular stick functions for cue and response/probe/target and a stretched/compressed stick function spanning the interval from cue to response/probe/target/decision. In particular, we estimated cue-locked responses that spanned from 200 ms pre-cue to 800 ms post-cue. The response/probe/target/decision response interval spanned from −800 to 200 ms. Each fixed-time response thus spanned 1000 ms, or 200 EEG sample points. The scaled-time component, as described earlier, was modelled as a single underlying component (set width in *X*) that spanned over multiple EEG durations (varying number of rows in *X*). Thus, our method required choosing how many scaled-time sample points to estimate (the width in *X*). For the production/perception tasks, we chose to estimate 330 scaled-time points, equivalent to the duration of the ‘medium’ interval. For the prediction task, we chose to estimate 200 scaled-time points, equivalent to the mean of the short and long conditions (700 ms, 1300 ms). For the decision-making task, we estimated 153 scaled-time points (roughly equivalent to the mean decision time). Unlike the conventional ERP approach, this analysis was conducted on the continuous EEG. To identify artifacts in the continuous EEG, we used the *basicrap* function from the ERPLAB [58] toolbox with a 150 μV threshold (2000 ms window, 1000 ms step size). A sample was flagged if it was ‘bad’ for any channel. Flagged samples were excluded from the GLM (samples removed from the EEG and rows removed from the design matrix). Additionally, we removed samples/rows associated with unusually fast or slow responses in the production task (less than 0.2 s or more than 5 s). On average, we removed 10.16 % of samples in the production task (95% CI [8.90, 11.42]), 3.75 % of samples in the perception task (95% CI [2.39, 5.10]), 5.57% of samples in the prediction task (95% CI [4.94, 6.20]), and 5.56% of samples in the decision-making task (95%CI [4.99, 6.12]).

To test for multicollinearity between the regressors we computed the variance inflation factor (VIF) for each regressor, i.e., at each timepoint in the estimated waveforms. This was done using the *uf_vif* function in the Unfold toolbox [25]. We were concerned about multicollinearity because the fixed-time and scaled-time components occurred over the same ‘real time’ durations. For example, in the production task the early and later parts of the scaled waveform always coincided with the start of the cue-locked and end of the response-locked responses, respectively. The overlap was not consistent, however; alignment between the fixed and scaled regressors was lessened due to distortions in the scaled stick function (see Supplementary Fig 1). As a result, the VIF was low (< 10) at nearly all points other than the start/end (Supplementary Fig 9). This was true in all tasks except for the temporal prediction task (VIFs > 10), as these tasks incorporated greater temporal variability across trials. We therefore expected the waveform estimates in the temporal prediction task to be noisier relative to the other tasks. We note that future studies could use VIF to evaluate the likelihood of successfully unmixing fixed-time and scaled-time components. Introducing elements of experimental design (such as increased interval variability across trials) could help to address concerns over multicollinearity.

To lessen the effect of multicollinearity and impose a smoothness constraint on our estimates, we used a first-derivative form of Tikhonov regularization [29]. Tikhonov regularization reframes the GLM solution as the minimization of:

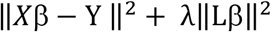

where *L* is the regularization operator and λ is the regularization parameter. In other words, we aimed to minimize a penalty term in addition to the usual residual. This has the solution

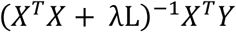

In our case, *L* approximated the first derivative as a scaled finite difference[59]:

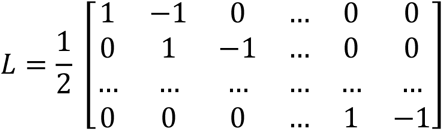

We then chose regularization parameters for each participant using 10-fold cross validation. Our goal here was to minimize the mean-squared error of the residual EEG at electrode FCz, our electrode of interest. The following λs were tested on each fold: 0.001, 0.01, 0, 1, 10, 100, 1000, 10000, 100000. An optimal λ was chosen for each participant corresponding to the parameter with the lowest mean mean-squared error across all folds. See Supplementary Table 7 for a summary.

For each task and participant, we computed the mean squared error (MSE) according to the model described above and – for comparison – a model with fixed-components only. To make the comparison fair, we only considered those timepoints for which the fixed components were active, e.g., from −200 ms to 800 ms relative to the cue and from −800 to 200 ms relative to the response.

#### Statistics

We quantified the amplitude of the scaled-time component by conducting a nonparametric statistical test of the scaled-time component according to the procedure outlined in [60]. After computing a single-sample *t*-statistic at each sample point and electrode, we identified clusters of points for which the *t*-value exceeded a critical threshold, corresponding to an alpha value of .001 for the production task, .01 for the perception task, and .05 for the prediction and decision-making tasks. Lower alpha values were used for the production and perception tasks to better isolate the effect; using an alpha value of .05 yielded longer windows of significance but did not change our results. Clusters were identified both spatially and temporally. For each electrode, we defined a cluster by identifying neighbouring electrodes according to a template available in the FieldTrip toolbox [61]. For each spatial cluster, we then identified temporal clusters for which the *t*-values of all the included electrodes exceeded the critical value. Within this common window we computed the ‘cluster mass’, defined as the spatial mean of the sum of the absolute values of the *t*-values within the temporal cluster. To determine whether the observed cluster masses exceeded what could occur by chance, we permuted the scaled components by randomly flipping (multiplying by −1) the entire waveform. We then computed and recorded the cluster masses for 1000 permuted waveforms. If more than one temporal cluster was found within a spatial cluster, only the maximum cluster mass was recorded. A value of zero was recorded if there were no clusters. We then labelled our observed cluster masses as ‘significant’ if they exceeded 95% of the maximum cluster masses of the permuted waveforms. Finally, we examined and reported the cluster extents and *p*-values for the clusters of maximum cluster mass: (P3, CP5, CP1) in the production task, (F3, Fz, FCz, C3, Cz) in the perception task, (FC1, FCz, FC2, C1, C2, CP1, CPz, CP2) in the prediction task, and (Cz, FC1, FCz, FC2, C1, C2, CP1, CPz, CP2) in the decision-making task.

#### Scaling Index

To validate the unmixing procedure, we regressed out either the scaled-time component or the fixed-time components from the EEG in each task and participant to create ‘fixed-only’ or ‘scaled-only’ data sets. We then quantified the amount of temporal scaling present in each task, participant, and data set (original, fixed-only, scaled only) using a similar procedure as [15]. Specifically, we constructed epochs spanning the intervals of interest (e.g., cue to response), then stretched or compressed each epoch to match a common duration (the longest duration in the interval timing tasks; the mean of the ‘late’ responses in the decision-making task, as defined above). For each task and participant, we averaged by condition (e.g., short, medium, long) to create conditional ERPs with a common duration, then computed a scaling index defined as the coefficient of determination. Specifically, we asked how well the ‘long’ waveform could be predicted by the ‘short’ waveform. If there was also a ‘medium’ waveform (the production/perception tasks) another coefficient of determination was computed, and the two coefficients were averaged. A larger scaling index can therefore be interpreted as a greater post-scaling similarity between conditions. Scaling indices in the fixed-only and scaled-only data sets were compared via paired-samples *t*-tests. For each *t*-test, we computed Cohen’s *d* as:

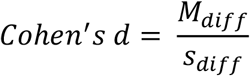

where *M_diff_* the mean difference between the scores being compared and s_diff_ is the standard deviation of the difference of the scores being compared [62]. Interestingly, the scaling index of the original signal appeared to be a mixture of the scaling indices of the fixed-only and scaled-only signals in all tasks except for the temporal prediction task (Fig 4). We interpreted this as further evidence that the unmixing procedure was less effective here due to multicollinearity.

#### PCA

To explore the link between the scaled-time component and behaviour, we examined the scaled-only data set described above – that is, the scaled-time regressors plus residuals. Only mid-frontal electrodes were considered: FC1, FCz, FC2, Cz, CP1, CPz, and CP2. We then constructed epochs starting at the cue and ending at the target interval (800 ms, 1650 ms, or 2500 ms). Epochs within each condition (short, medium, long) were further grouped into three equal-sized response-time bins (early, on time, late) and averaged for each electrode and participant. We then conducted a PCA for each condition (short, medium, long) and participant. See Supplementary Table 5 for amount of variance explained by PC1 and PC2. To visualize the effect of PC2, we computed the mean PC2 across all participants. We then added more or less of the mean PC2 to the mean PC1 projection and applied a 25-point moving-mean window for visualization purposes (Supplementary Fig 4). In order to choose a reasonable range of PC2 scores, we examined the average minimum and maximum PC2 score for each participant and condition (short, medium, long). The PC2 score ranges were − 21 to 15 (short), −41 to 38 (medium), and −40 to 55 (long). To assess the relationship between PC2 score and behaviour, we binned PC2 scores according to our response time bins (early, on time, late) and collapsed across conditions (short, medium, long). This gave us as single mean PC2 score for each participant and response time bin (early, on time, late), which we analyzed using a two-sided repeated-measures ANOVA (Fig 5d) after verifying the assumption of normality using the Shapiro-Wilk test. Two different effect sizes, η*_p_*^2^ and η*_g_*^2^, were computed, according to:

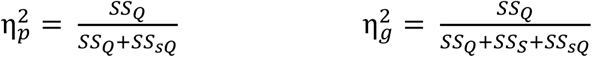

where *SS*_Q_ is the sum of squares of the quantile effect (early, on time, late), *SS_sQ_* is the error sum of squares of the quantile effect, and *SS_S_* is the sum of squares between subjects [63].

We then replicated the PCA procedure for the decision-making task using an epoch extending 800 ms from the cue at a central electrode cluster (FC3, FC1, FC2, FC4, C3, C1, Cz, C2, C4, CP3, CP1, CP2, CP4, P3, P1, Pz, P2, and P4). Note that the assumption of normality was violated for ‘early’ responses in the decision-making task. However, as repeated-measures ANOVA is robust to violations of normality, no statistical correction was made.

#### Data Availability

Raw data for the production and perception tasks is available at https://doi.org/10.18112/openneuro.ds004200.v1.0.0. Raw data for the prediction task is available at https://doi.org/10.5061/dryad.5vb8h. Raw data for the decision-making task is available at https://doi.org/10.18112/openneuro.ds002734.v1.0.2.

#### Code Availability

Simulation and analysis scripts are available at https://github.com/chassall/temporalscaling.

