## Supplementary Material for "Temporal scaling of human scalp-recorded potentials"

Supplementary Table 1
*Probe times in the perception task (in seconds), for each condition (short, medium, long)*

| | Very early | Early | On time | Late | Very late |
|---|---|---|---|---|---|
| Short | 0.512 | 0.640 | 0.800 | 1.00 | 1.250 |
| Medium | 1.056 | 1.320 | 1.650 | 2.0625 | 2.5781 |
| Long | 1.6 | 2 | 2.5 | 3.125 | 3.9062 |

Supplementary Table 2
*Mean response times in the production task for each condition (short, medium, long)*

| | Mean (s) | 95% CI |
|---|---|---|
| Short | 0.807 | [0.770, 0.844] |
| Medium | 1.634 | [1.565, 1.703] |
| Long | 2.319 | [2.239, 2.398] |

Supplementary Table 3
*Mean proportion of "yes" response in the perception task for each condition (short, medium, long) and probe time (v. early, early, on time, late, v. late)*

| | Short | | Medium | | Long | |
|---|---|---|---|---|---|---|
| | Mean (%) | 95% CI | Mean (%) | 95% CI | Mean (%) | 95% CI |
| Very Early | 7.92 | [2.19, 13.64] | 10.83 | [4.62, 17.05] | 12.92 | [5.93, 19.90] |
| Early | 38.75 | [30.14, 47.36] | 40.00 | [31.55, 48.45] | 39.17 | [27.02, 51.31] |
| On Time | 84.58 | [78.22, 90.94] | 69.17 | [60.96, 77.38] | 57.50 | [49.92, 65.08] |
| Late | 40.83 | [28.57, 53.10] | 33.13 | [25.06, 41.19] | 23.13 | [16.72, 29.53] |
| Very Late | 12.50 | [4.35, 20.65] | 6.88 | [2.52, 11.23] | 5.83 | [2.98, 8.69] |

Supplementary Table 4
*Mean response time in the prediction ask for each condition (rhythmic, repeated) and interval (short, long).*

| | Rhythmic | | Repeated | |
|---|---|---|---|---|
| | Mean (s) | 95% CI | Mean (s) | 95% CI |
| Short | 0.228 | [0.216, 0.241] | 0.229 | [0.212, 0.246] |
| Long | 0.228 | [0.215, 0.241] | 0.226 | [0.213, 0.239] |

Supplementary Table 5
*PCA variance explained by PC1 and PC2 in the production task (short, medium, and long conditions) and in the decision-making task.*

| | PC1 | | PC2 | |
|---|---|---|---|---|
| | Variance (%) | 95% CI | Variance (%) | 95% CI |
| Production (Short) | 89.20 | [85.63, 92.77] | 4.54 | [3.05, 6.02] |
| Production (Medium) | 88.84 | [85.28, 92.39] | 5.03 | [3.21, 6.84] |
| Production (Long) | 89.68 | [86.55, 92.80] | 4.23 | [2.91, 5.55] |
| Decision-Making | 78.58 | [74.06, 83.11] | 9.79 | [7.89, 11.69] |

Supplementary Table 6
*Mean PC2 score by quantile in the interval production and decision-making tasks.*

| | Early | | On Time | | Late | |
|---|---|---|---|---|---|---|
| | Mean | 95% CI | Mean | 95% CI | Mean | 95% CI |
| Production | 11.56 | [1.24, 21.87] | -1.06 | [-8.65, 6.53] | -11.50 | [-21.50, -1.51] |
| Decision Making | 7.55 | [2.70, 12.41] | 2.36 | [-0.24, 4.97] | -2.30 | [-7.70, 3.10] |

Supplementary Table 7

*Optimal regularization parameters in each task by number of participants*.

| | Optimal λ (participant count) | | | | | | | | | | |
|---|---|---|---|---|---|---|---|---|---|---|---|
| | $10^{-3}$ | $10^{-2}$ | $10^{-1}$ | $10^{0}$ | $10^{1}$ | $10^{2}$ | $10^{3}$ | $10^{4}$ | $10^{5}$ | $10^{6}$ | $10^{7}$ |
| Production | 0 | 0 | 0 | 0 | 0 | 0 | 2 | 18 | 0 | 0 | 0 |
| Perception | 0 | 0 | 0 | 0 | 0 | 0 | 0 | 20 | 0 | 0 | 0 |
| Prediction | 0 | 0 | 0 | 0 | 0 | 7 | 12 | 0 | 0 | 0 | 0 |
| Decision-Making | 0 | 0 | 0 | 0 | 0 | 0 | 0 | 18 | 0 | 0 | 0 |

Supplementary Table 8

*Mean squared errors for the full model and a fixed-only model (*x $10^7$ $\mu V^2$)

| | Full | | Fixed-Only | | | | |
|---|---|---|---|---|---|---|---|
| | MSE | 95% CI | MSE | 95% CI | *t* | *p* | Cohen's *d* |
| Production | 1.7005 | [1.1631, 2.2380] | 1.7088 | [1.1697, 2.2478] | -3.97 | <.001 | -0.89 |
| Perception | 1.9175 | [1.3233, 2.5116] | 1.9247 | [1.3286, 2.5209] | -5.09 | <.001 | -1.14 |
| Prediction | 1.4987 | [1.1606, 1.8369] | 1.5124 | [1.1729, 1.8520] | -4.90 | <.001 | -1.12 |
| Decision-Making | 0.5112 | [0.4697, 0.5526] | 0.5124 | [0.4709, 0.5539] | -7.77 | <.001 | -1.83 |

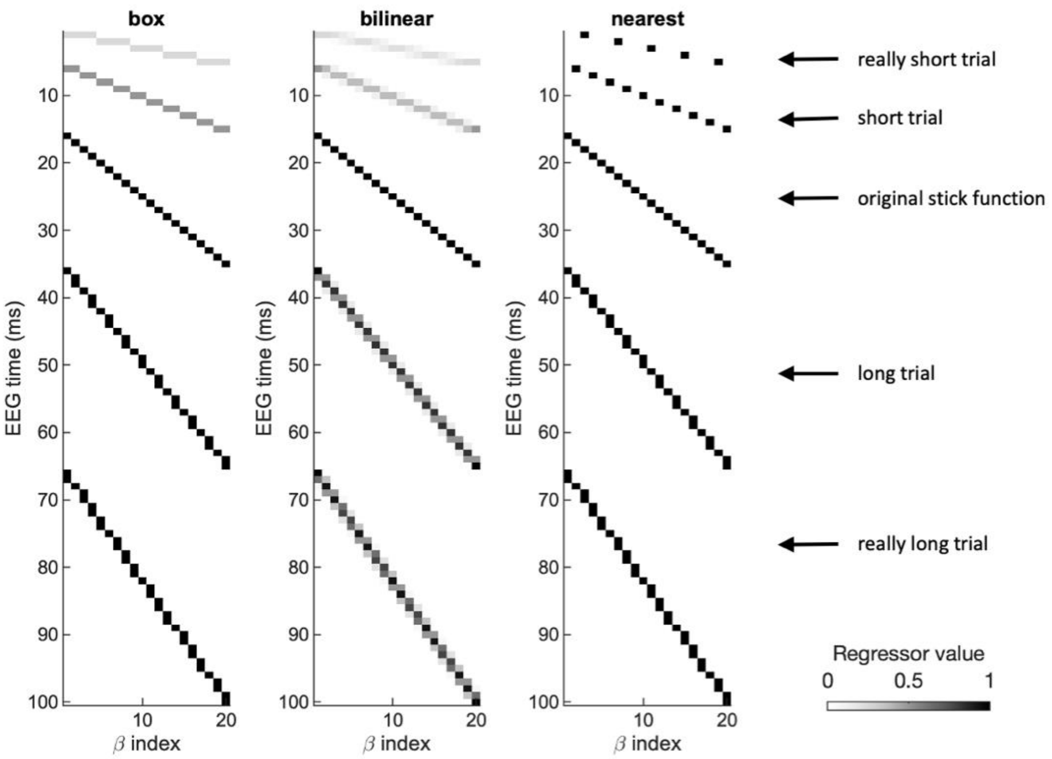


**Supplementary Fig 1. Compressing/stretching a stick function to model scaled-time components.** A stick function was compressed or stretched to model a common neural response that unfolded over varying timescales. The function was interpolated using a box-shaped kernel – gaps in the stretched function were 'filled in' with ones, whereas vertically-adjacent values (ones and zeroes) were averaged in the compressed function. Two other interpolation methods are shown (linear interpolation and a nearest-neighbours approach). For the purpose of illustration, only 20 beta values are plotted. The actual number of scaled-time beta values was 330 in the production and perception tasks, and 200 in the prediction task.

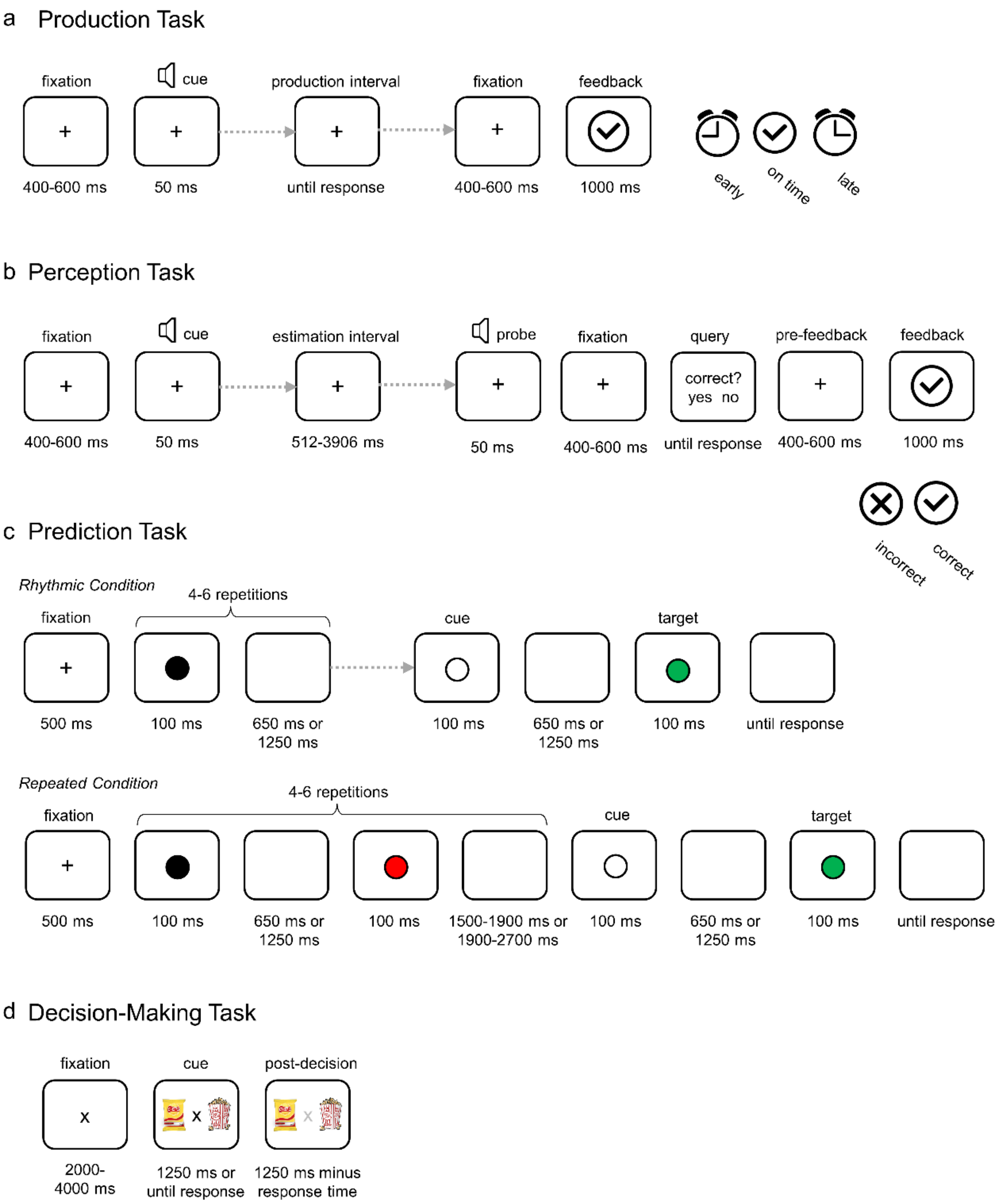


**Supplementary Fig 2. Timing details for each task.** (a) In the production task participants reproduced a short, medium, or long temporal interval. Visual feedback indicated whether participants were early, on time or late. (b) In the perception task participants decided whether or not a computer-produced interval (short, medium, long) was on time. Visual feedback indicated the correctness of the decision (a checkmark or an ‘x’). Each block of (a) and (b) was preceded by a metronome indicating the target interval. (c) Two out of four conditions of the prediction task were analyzed. The conditioned differed in the rhythmicity of the temporal cue. In the rhythmic condition, the cue was a visual stimulus that flickered at a constant rate. In the repeated condition, the cue was a pair of stimuli separated by a constant duration but followed by a variable jitter period. In each condition, the warning cue then appeared followed by the target stimulus. Participants were instructed to respond to the target stimulus as quickly as possible. (d) In the decision-making task, participants were presented with images of two snack foods and asked to select their preferred option. Decision time varied up to a maximum of 1250 ms (the trial time limit).

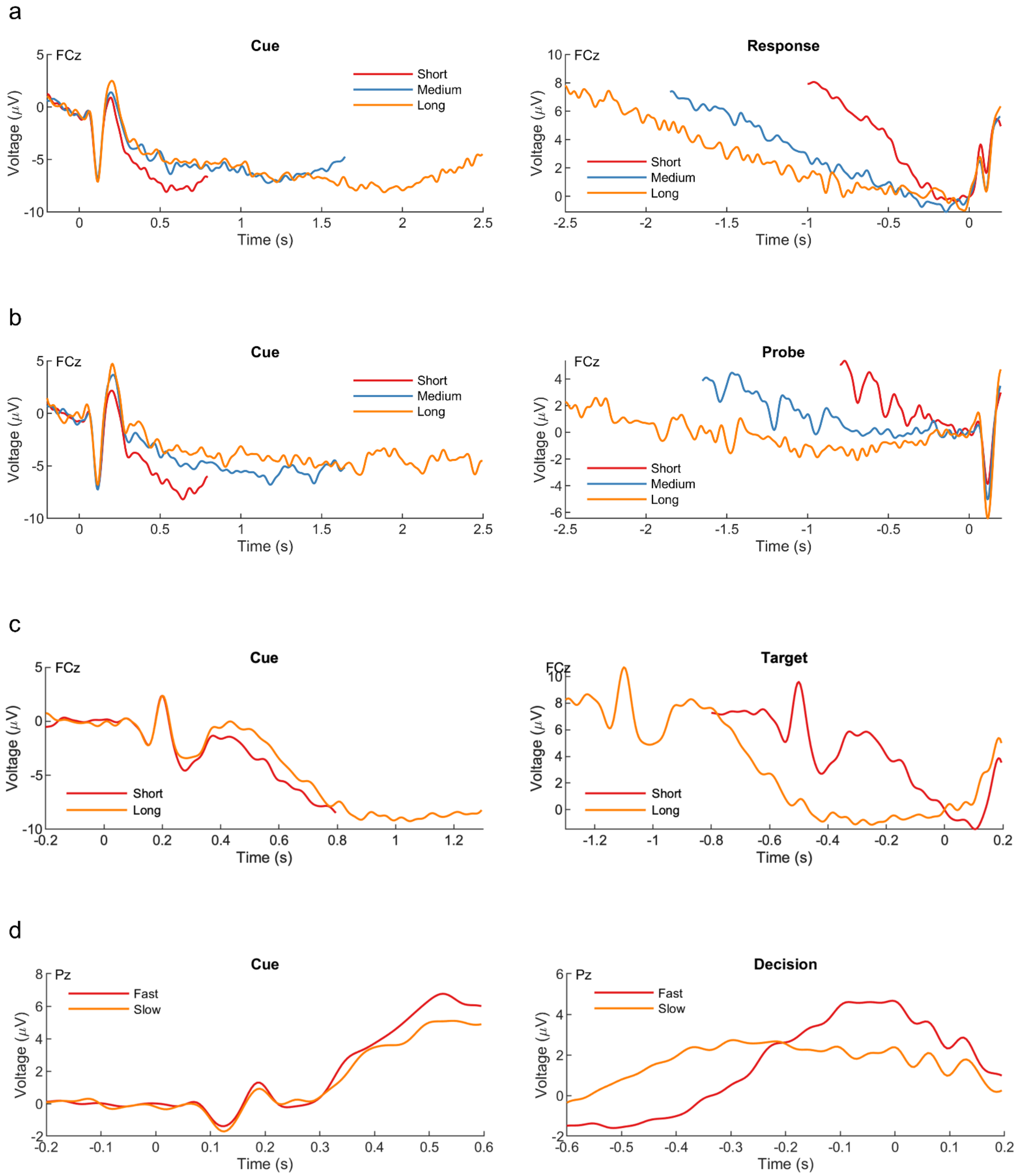


**Supplementary Fig 3. A conventional ERP analysis reveals ramping activity during the delay period.** Differences were apparent in the cue-locked and response-locked activity in the (a) production task, (b) perception task, and (c) prediction task.

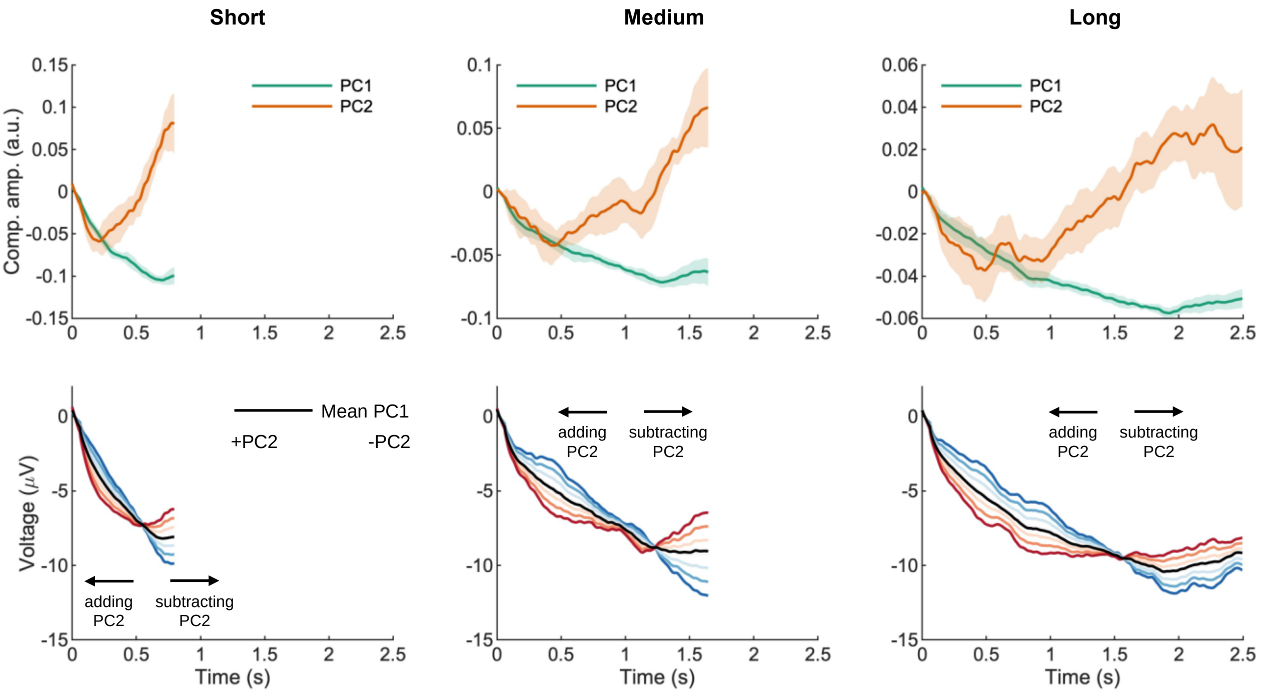


**Supplementary Fig 4. Principal components capture signal amplitude and latency in the temporal production task.** Cue-locked EEG over a central-parietal cluster (P3, CP5, CP1) was grouped by response time (early, on time, or late), averaged, and stacked for each target interval (short, medium, or long). A separate PCA was run for each target interval and participant. In each condition, the first two principal components for each target interval represent the amplitude (PC1) and first derivative (PC2) of the time-scaled component (top). Adding or subtracting different amounts of PC2 to PC1 shifted the peak earlier or later in time (bottom). Only electrode P3 is shown (the cluster centre).

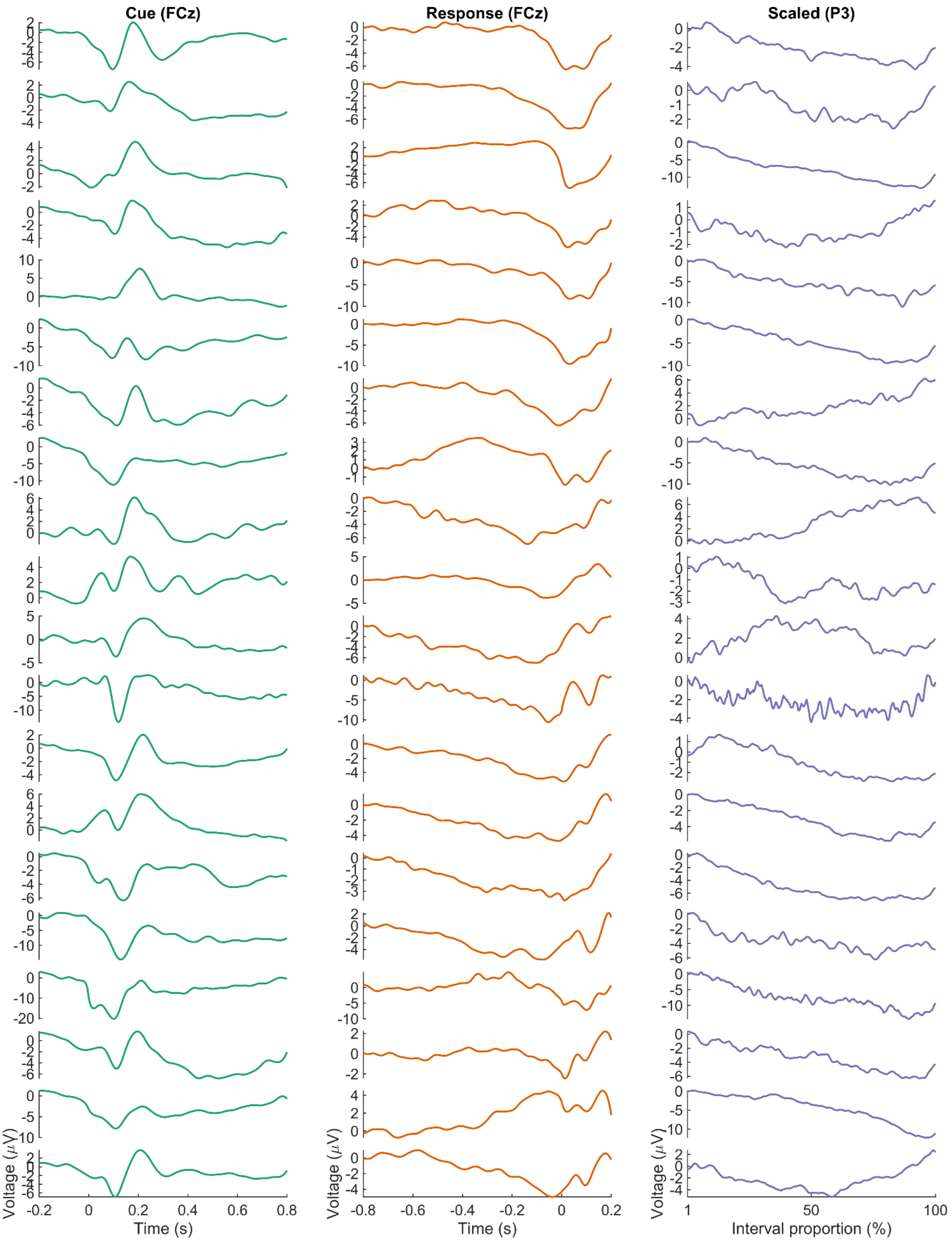


**Supplementary Fig 5. Every participant's regression-ERP in the production task.**

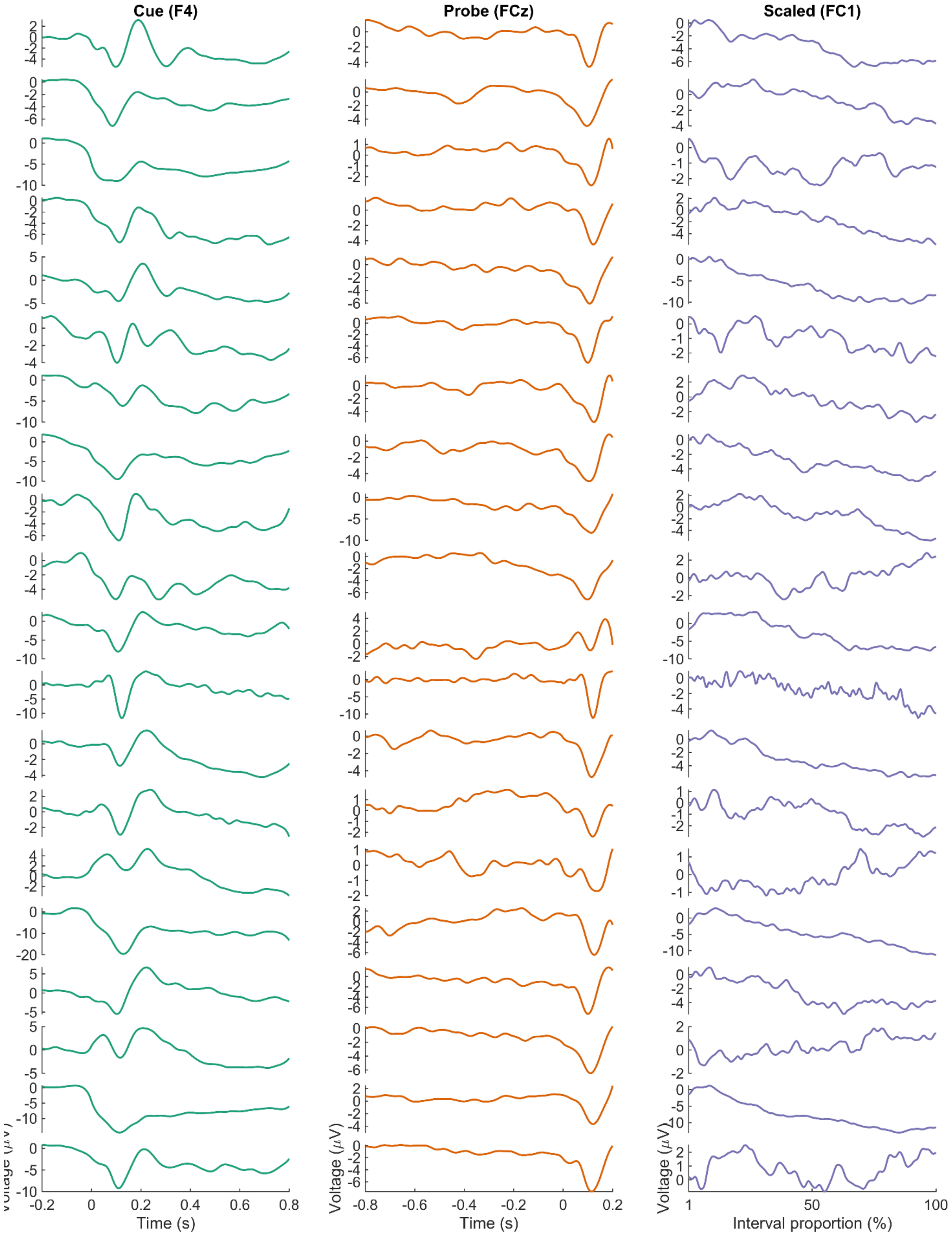


**Supplementary Fig 6. Every participant's regression-ERP in the perception task.**

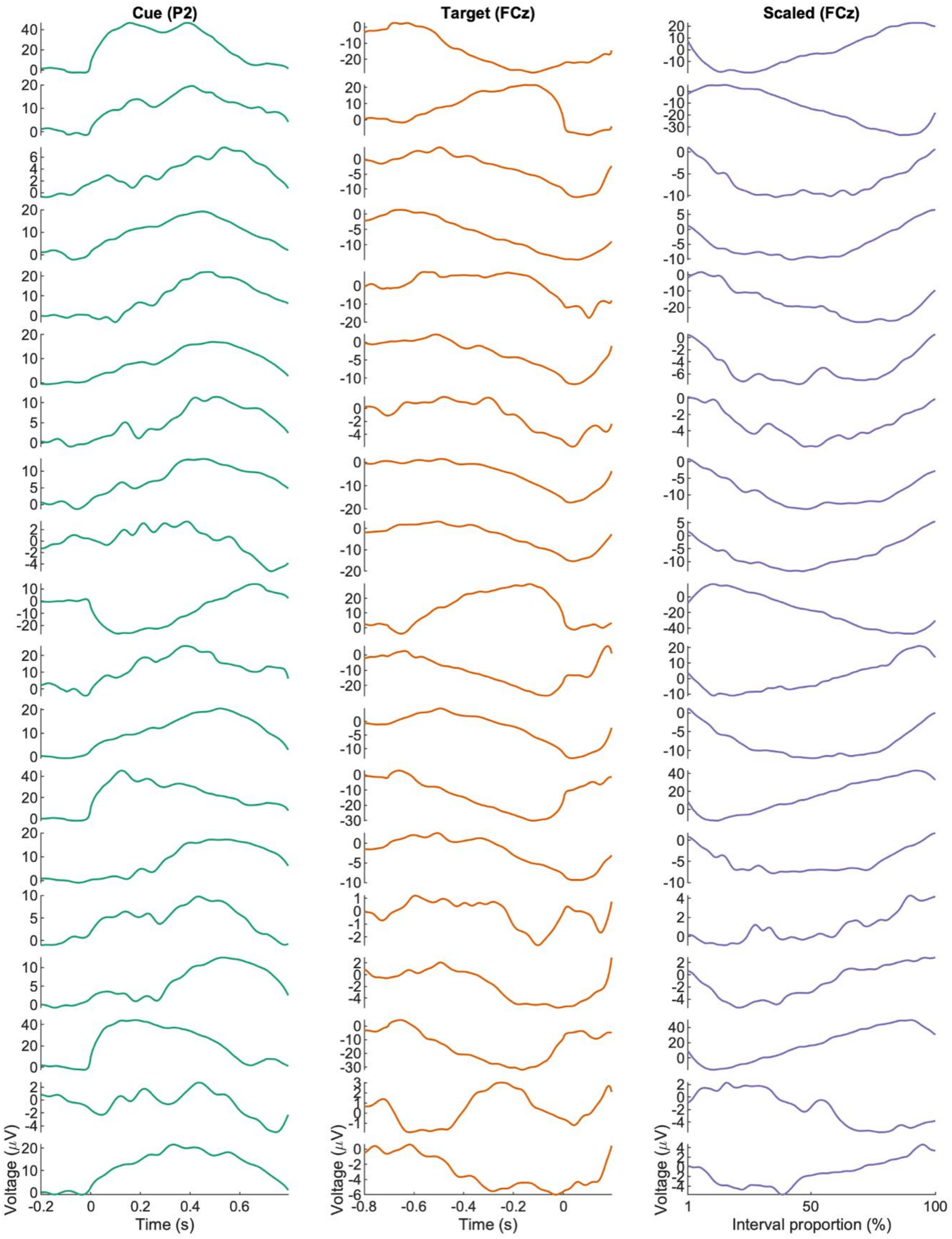


**Supplementary Fig 7. Every participant's regression-ERP in the prediction task.**

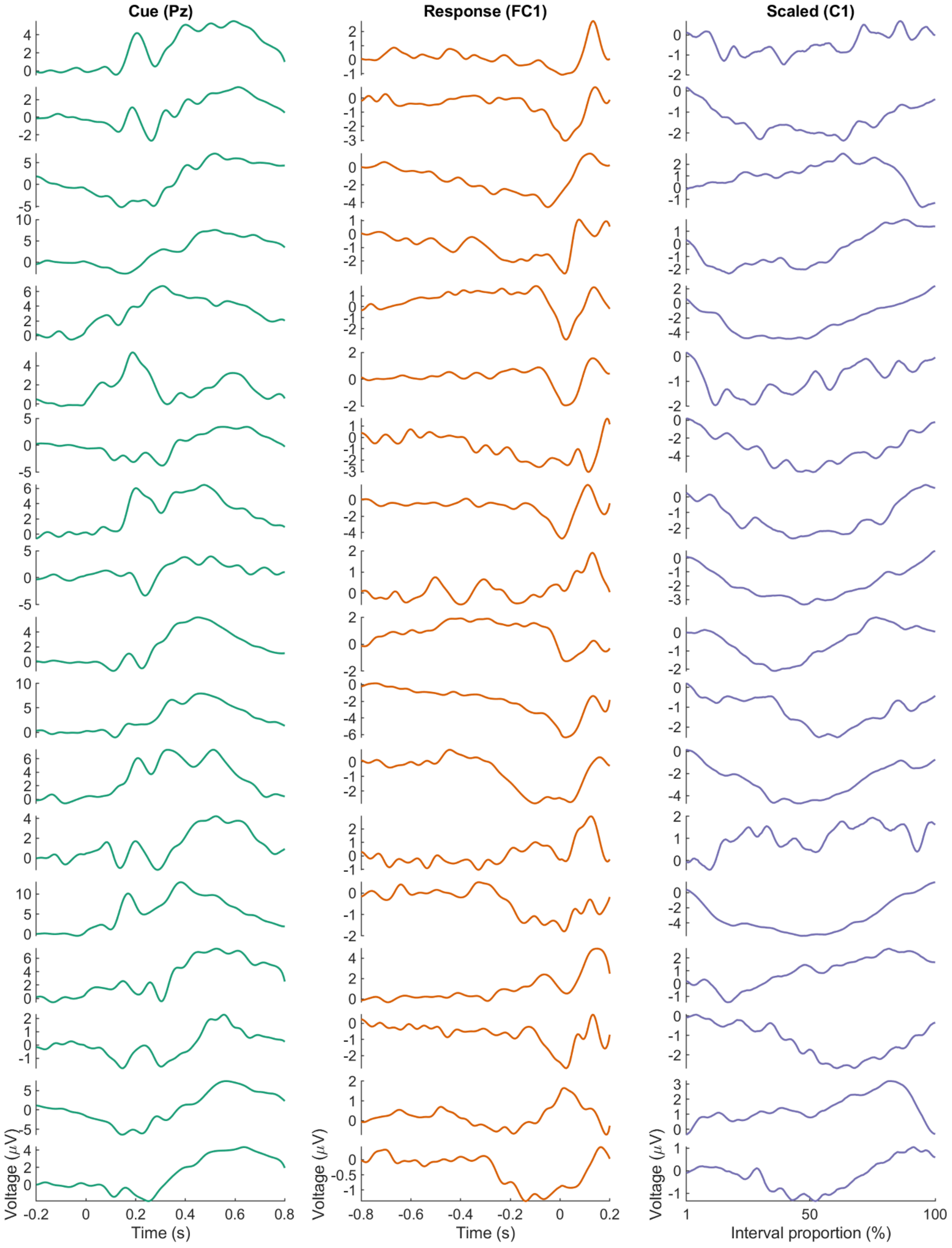


**Supplementary Fig 8. Every participant's regression-ERP in the decision-making task.**

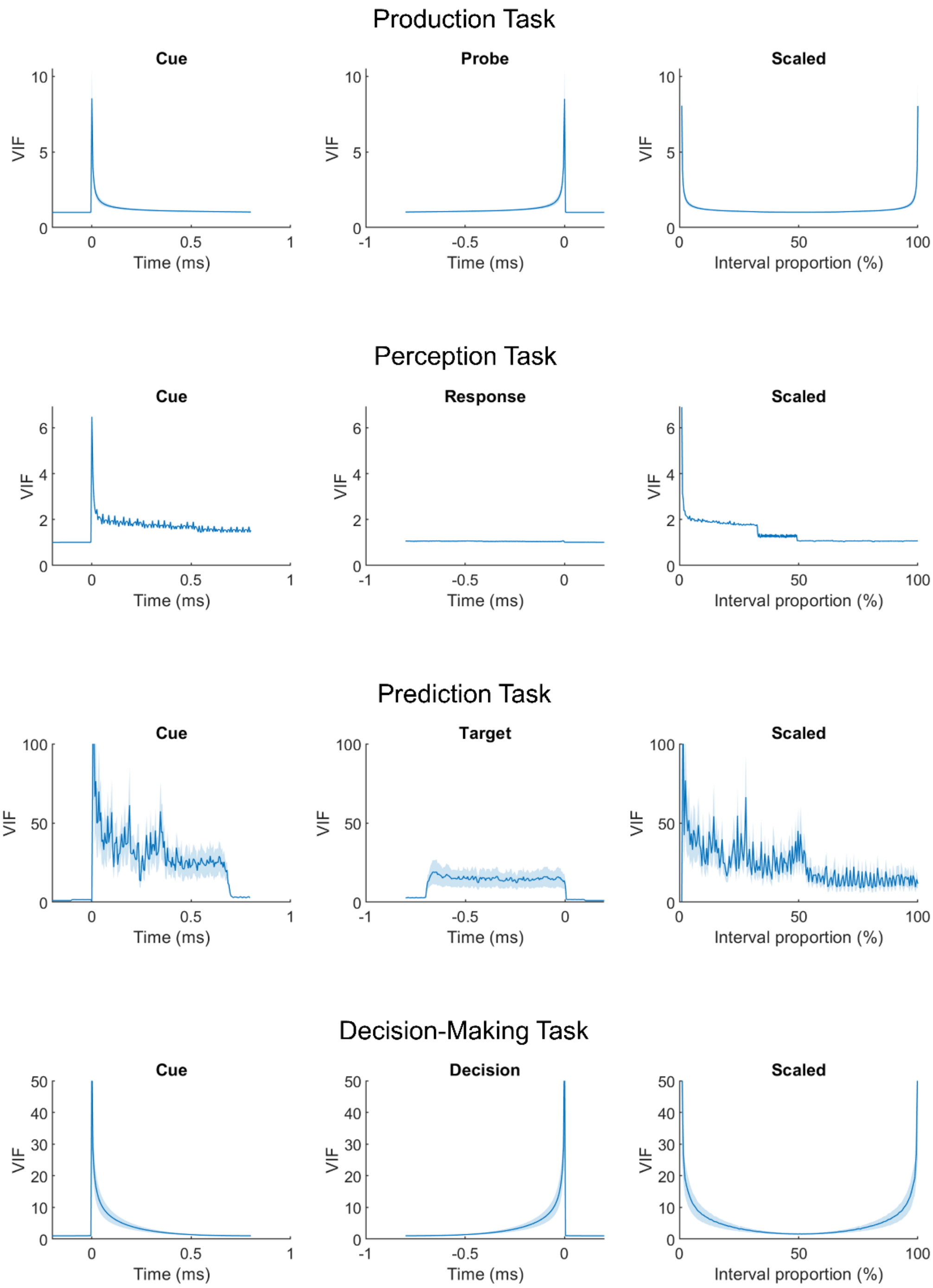


**Supplementary Fig 9. Variance inflation factor (VIF) in each task.**
